# No evidence for functional distinctions across fronto-temporal language regions in their temporal receptive windows

**DOI:** 10.1101/712372

**Authors:** Idan A. Blank, Evelina Fedorenko

**Author notes:** **Corresponding Author:** Idan A. Blank.

## Abstract

The “core language network” consists of left temporal and frontal regions that are selectively engaged in linguistic processing. Whereas the functional differences across these regions have long been debated, many accounts propose distinctions in terms of representational grain-size—e.g., words *vs.* sentences—or processing time-scale, i.e., operating on local linguistic features *vs.* larger spans of input. Indeed, the topography of language regions appears to overlap with a cortical hierarchy reported by Lerner et al. (2011) wherein mid-posterior temporal regions are sensitive to low-level features of speech, surrounding areas—to word-level information, and inferior frontal areas—to sentence-level information and beyond. However, the correspondence between the language network and this hierarchy of “temporal receptive windows” (TRWs) is difficult to establish because the precise anatomical locations of language regions vary across individuals. To directly test this correspondence, we first identified language regions in each participant with a task-based localizer, which confers high functional resolution to the study of TRWs (traditionally based on stereotactic coordinates); then, we characterized regional TRWs with the naturalistic story listening paradigm of Lerner et al. (2011), which augments task-based characterizations of the language network by more closely resembling comprehension “in the wild”. We find no region-by-TRW interactions across temporal and inferior frontal regions, which are all sensitive to both word-level and sentence-level information. Therefore, the language network as a whole constitutes a unique stage of information integration within a broader cortical hierarchy.

**Highlights:**

- Language regions are identified with task-based, participant-specific localization.
- A progressively scrambled naturalistic story probes regional processing timescales.
- Widespread sensitivity to scrambling at the timescales of both words and sentences.
- No processing timescale distinctions across temporal and inferior-frontal regions.
- These regions all occupy a common, unique stage in a broader processing hierarchy.

## 1. Introduction

Language comprehension engages a cortical network of frontal and temporal brain regions, primarily in the left hemisphere (Binder et al., 1997; Bates et al., 2003; Fedorenko et al., 2010; Menenti et al., 2011). There is ample evidence that this “core language network” is language-selective and is not recruited by other mental processes (Fedorenko and Varley, 2016; see also Pritchett et al., 2018; Ivanova et al., 2019; Jouravlev et al., 2019), indicating that it either employs cognitively unique representational formats and/or implements algorithms distinct from those recruited in other cognitive domains. Nonetheless, the functional architecture of this network—i.e., the division of linguistic labor among its constituent regions—remains highly debated. On the one hand, some neuroimaging studies have suggested that different linguistic processes are localized to focal and distinct subsets of this network (e.g., Stowe et al., 1998; Vandenberghe et al., 2002; Bornkessel et al., 2005; Humphries et al., 2006; Caplan et al., 2008; Snijders et al., 2009; Meltzer et al., 2010; Pallier et al., 2011; Brennan et al., 2012; Goucha and Friederici, 2015; Zhang and Pylkkänen, 2015; Kandylaki et al., 2016; Frank and Willems, 2017; Wilson et al., 2018; Bhattasali et al., 2019). On the other hand, other studies have reported that such processes are widely distributed and spatially overlapping (e.g., Keller et al., 2001; Vigneau et al., 2006; Fedorenko et al., 2012b; Bautista and Wilson, 2016; Blank et al., 2016; Fedorenko et al., 2018; Siegelman et al., 2019). Similar conundrums are also characteristic of the neuropsychological (patient) literature (Caplan et al., 1996; Dick et al., 2001; Bates et al., 2003; Wilson and Saygin, 2004; Grodzinsky and Santi, 2008; Tyler et al., 2011; Duffau et al., 2014; Mesulam et al., 2015; Mirman et al., 2015; Fridriksson et al., 2018; Matchin and Hickok, 2019).

Proposals for the functional architecture of the core language network vary substantially from one another in the theoretical constructs posited and the mapping of those constructs onto brain regions (for examples, see Friederici, 2002; Hickok and Poeppel, 2004; Ullman, 2004; Grodzinsky and Friederici, 2006; Hickok and Poeppel, 2007; Bornkessel-Schlesewsky and Schlesewsky, 2009; Friederici, 2011, 2012; Poeppel et al., 2012; Price, 2012; Bornkessel-Schlesewsky and Schlesewsky, 2013; Hagoort, 2013). Such differences notwithstanding, the majority of accounts share a common, fundamental hypothesis: different language regions integrate incoming linguistic input at distinct, progressively longer timescales. This hypothesis may take different forms: in some accounts, linguistic representations of different grain sizes (e.g., phonemes, morphemes, words, phrases and sentences) are respectively mapped onto distinct regions; in other accounts, some regions function as mental lexicons (“memory”) that store smaller linguistic units, whereas other regions combine these units into larger structures (“online processing”). Yet all forms of this hypothesis, while considerably varying in critical details, agree that a functional dissociation among language regions would manifest as differences in their respective timescales for processing and integration.

A brain region’s integration time scale constrains the amount of preceding context that influences the processing of current input. A relatively short integration timescale entails that the incoming signal is integrated with its local context, with more global context exerting less influence, whereas a longer integration timescale entails sensitivity to broader contexts extending farther into the past. This context sensitivity governs how closely a region could track linguistic input that deviates from well-formedness: a brain region with a short integration timescale (e.g., on the order of single morphemes) should reliably track any locally well-formed input even in the face of coarser, global disorder (morphemes can be extracted even from ungrammatical sequences of unrelated words); but a region with a longer integration timescale (e.g., on the order of phrases) could not reliably track such locally-intact-yet-globally-incoherent input (phrases would be difficult to establish in such sequences). Therefore, a straightforward prediction that follows from the “hierarchy of processing timescales” hypothesis is that different language regions should exhibit distinct patterns of input tracking when well-formedness deteriorates from coarser, more global disruptions to finer, more local violations.

Indeed, such a response profile consistent with a hierarchy of integration timescales has been reported in a set of left temporal and frontal areas, whose topography appears consistent with the core language network (Lerner et al., 2011). Specifically, Lerner and colleagues played participants a naturalistic spoken story along with several, increasingly scrambled versions of it: a list of unordered paragraphs, a list of unordered sentences, a list of unordered words, and the audio recording played in reverse. As participants listened to each of these stimuli in a fMRI session, voxel-wise fluctuations in BOLD signal were recorded and the reliability of their input tracking was then evaluated. Lerner and colleagues reasoned that if neurons in a given voxel were able to reliably track and integrate a certain stimulus, then the resulting signal fluctuations would be stimulus-locked and, thus, similar across individuals; in contrast, untracked input would elicit fluctuations that would not be reliably related to the stimulus and, thus, would differ across individuals. Therefore, the authors computed for each voxel and stimulus an inter-subject correlation (ISC; Hasson et al., 2004) of BOLD signal fluctuations. Their novel approach revealed a hierarchy of integration timescales (or “temporal receptive windows”; TRWs) extending from mid-temporal regions both rostrally and caudally along the temporal lobe and on to inferior frontal regions:

Mid-temporal regions early in the hierarchy reliably tracked all stimuli including the reverse audio and the word list, which indicated a short integration timescale. Next, a little more posteriorly and anteriorly, temporal regions tracked all stimuli except for the reverse audio, indicative of sensitivity to sub-word (e.g., morpheme) or word-level information (a short TRW). Further posterior and anterior temporal regions could only track lists of sentences or paragraphs (but not words), indicative of sensitivity to phrase- or sentence-level information. And, finally, some inferior frontal regions exhibited this same pattern of sensitivity to phrase/sentence information, with yet others reliably tracking only paragraph (but not sentence) lists, indicative of sensitivity to information above the sentence level (a long TRW).

This hierarchy of integration timescales is an appealing organizing principle of the core language network (DeWitt and Rauschecker, 2012; Bornkessel-Schlesewsky et al., 2015; Hasson et al., 2015; Chen et al., 2016; Baldassano et al., 2017; Yeshurun et al., 2017a; Sheng et al., 2018). Nevertheless, there are several reasons to question the putative correspondence between this hierarchy and the set of language-selective cortical regions. The first issue is neurobiological: the process of TRW characterization described above is carried out on a voxel-by-voxel basis and, hence, crucially relies on the assumption that a given voxel houses the same functional circuits across individuals, but this assumption is demonstrably invalid. There is significant inter-individual variability in the mapping of functions onto macro-anatomy (Duffau, 2017), especially in the frontal and temporal lobes, both associative areas that are functionally heterogeneous (Jones and Powell, 1970; Gloor, 1997; Amunts et al., 1999; Tomaiuolo et al., 1999; Wise et al., 2001; Chein et al., 2002; Juch et al., 2005; Fedorenko et al., 2012a; Deen et al., 2015). Therefore, the same stereotactic coordinate may be part of the core language network in one brain but part of a functionally distinct network in another (Poldrack, 2006; Fischl et al., 2008; Frost and Goebel, 2012; Tahmasebi et al., 2012), in which case voxel-based inter-subject correlations in BOLD signal fluctuations cannot be interpreted as markers of input tracking by a specific functional circuit.

The second issue is statistical. Even if different language regions showed evidence of differing integration timescales at the descriptive level, direct statistical comparisons across their response profiles would be required in order to establish that they are functionally distinct. For instance, when the tracking of an unordered list of unrelated words is significant in one region but not in another, the difference between these two regions might itself still be non-significant (Nieuwenhuis et al., 2011). In other words, a region-by-stimulus interaction test is a crucial piece of statistical evidence in support of a hierarchy of integration timescales within the core language network, but such a test has hitherto been missing.

The third issue pertains to psycholinguistic theory and data. Although there is little doubt that comprehension proceeds via a cascaded integration of input along increasingly longer timescales (constructing larger meaningful units out of smaller ones; see, e.g., Christiansen and Chater, 2016), different stages of this process need not rely on qualitatively distinct mental structures or memory stores. Instead, language processing operates over a continuum of representations that straddle the traditionally postulated boundaries between sounds and words (Farmer et al., 2006; Bradlow and Bent, 2008; Maye et al., 2008; Trude and Brown-Schmidt, 2012; Schmidtke et al., 2014) or between words and larger constructions and combinatorial rules (Clifton et al., 1984; MacDonald et al., 1994; Trueswell et al., 1994; Garnsey et al., 1997; Traxler et al., 2002; Reali and Christiansen, 2007; Gennari and MacDonald, 2008) (see also Joshi et al., 1975; Schabes et al., 1988; Goldberg, 1995; Bybee, 1998; Jackendoff, 2002; Culicover and Jackendoff, 2005; Wray, 2005; Bybee, 2010; Snider and Arnon, 2012; Jackendoff, 2007; Langacker, 2008; Christiansen and Arnon, 2017). By extension, linguistic representations of different grain sizes need not be spatially segregated in the cortex across distinct regions and rely on dissociable neural resources.

Therefore, the current study directly tested for a functional dissociation among core language regions in terms of their temporal receptive windows. To this end, and to address the methodological issues discussed above, we synergistically combined two neuroimaging paradigms with complementary strengths: a traditional, task-based design and a naturalistic, task-free design. First, we used a well validated localizer task to identify regions of the core language network individually in each participant (Fedorenko et al., 2010; Julian et al., 2012). This approach allowed us to establish correspondence across brains based on functional response profiles (Saxe et al., 2006) rather than stereotaxic coordinates in a common space, thus augmenting the common voxel-based methodologies for studying temporal receptive windows. Then, we characterized the temporal receptive windows in each language region by using the same naturalistic story and its scrambled versions from Lerner et al. (2011). This paradigm broadly samples the space of computations and representations engaged during comprehension and, thus, tests the “hierarchy of processing timescales” hypothesis in its most general formulation, untied to more detailed theoretical commitments. It therefore augments task-based paradigms, which rely on carefully crafted materials and tasks that isolate particular mental processes tied to specific constructs. Finally, we computed inter-subject correlations for each stimulus in each functionally localized language region, and tested for region-by-stimulus interaction to directly compare the resulting temporal receptive windows across the network. In sum, our combined approach both (i) enjoys the increased ecological validity of naturalistic, “task-free” neuroimaging paradigms that mimic comprehension “in the wild” rather than “in the lab”; and (ii) guarantees the functional interpretability of the studied regions by harnessing a participant-specific localizer task instead of relying on precarious “reverse inference” from anatomy back to function (Poldrack, 2006, 2011).

## 2. Materials and Methods

### 2.1. Participants

Nineteen participants (12 females) between the ages of 18 and 47 (median=22), recruited from the MIT student body and the surrounding community, were paid for participation. All participants were native English speakers, had normal hearing, and gave informed consent in accordance with the requirements of MIT’s Committee on the Use of Humans as Experimental Subjects (COUHES). One more participant was originally scanned but excluded from analysis due to poor behavioral performance and neuroimaging data quality.

### 2.2. Design, materials and procedure

Each participant performed the language localizer task (Fedorenko et al., 2010) and, for the critical experiment, listened to five versions of a narrated story (cf. Lerner et al., 2011, where different subsets of the sample listened to different subsets of the stimulus set). The localizer and critical experiment were run either in the same scanning session (13 participants) or in two separate sessions (6 participants, who have previously performed the localizer task while participating in other studies) (see Mahowald and Fedorenko, 2016 for evidence of high stability of language localizer activations over time). In each session, participants performed a few other, unrelated tasks, with scanning sessions lasting 90-120min.

#### 2.2.1. Language localizer task

Regions in the core language network were localized using a passive reading task that contrasted sentences (e.g., DIANE VISITED HER MOTHER IN EUROPE BUT COULD NOT STAY FOR LONG) and lists of unconnected, pronounceable nonwords (e.g., MARSANORES CHENDEN LIPE FOR LANTIE THE THAT FALPS COMTERED THE PRINE IN) (Fedorenko et al., 2010). Each stimulus consisted of 12 words/nonwords, presented at the center of the screen one word/nonword at a time at a rate of 450ms per word/nonword. Each trial began with 100ms of fixation and ended with an icon instructing participants to press a button, presented for 400ms and followed by 100ms of fixation, for a total trial duration of 6s. The button-press task was included to help participants remain alert and focused throughout the run. Trials were presented in a standard blocked design with a counterbalanced order across two runs. Each block, consisting of 3 trials, lasted 18s. Fixation blocks were evenly distributed throughout the run and lasted 14s. Each run consisted of 8 blocks per condition and 5 fixation blocks, lasting a total of 358s. (A version of this localizer is available for download from https://evlab.mit.edu/funcloc/download-paradigms)

The *sentences > nonwords* contrast targets high-level aspects of language, to the exclusion of perceptual (speech / reading) and motor-articulatory processes (for a discussion, see Fedorenko and Thompson-Schill, 2014; Fedorenko, in press). We chose to use this particular localizer contrast for compatibility purposes with other past and ongoing experiments in the Fedorenko lab. For the current study, the main requirements from the localizer contrast were that it neither be under-inclusive nor over-inclusive. Below, we address each requirement in turn.

First, to avoid under-inclusiveness, the contrast should identify regions engaged in a variety of high-level linguistic processes, from ones that might depend on local information integration (e.g., single-word processing) to those that might depend on more global integration (e.g., processing of multi-word constructions, online composition). Because sentences differ from nonword lists in requiring processing at both the word level and above, the contrast’s content validity is appropriate. Moreover, this localizer contrast has been extensively validated over the past decade, and it identifies regions that all exhibit sensitivity to word-, phrase-, and sentence-level semantic and syntactic processing (Fedorenko et al., 2010; Fedorenko et al., 2012b; Blank et al., 2016; Mollica et al., 2018), which are the levels we focus on as described in the Results section. For instance, the regions identified with this localizer all exhibit reliable effects in narrower contrasts, e.g., sentences *vs.* word lists; sentences *vs.* “Jabberwocky” sentences (where content words have been replaced with nonwords); word lists *vs.* nonword lists; and “Jabberwocky” sentences *vs.* nonword lists (similar patterns obtain in electrocorticographic data with high temporal resolution: Fedorenko et al., 2016). In addition, contrasts that are broader than *sentences > nonwords* and that do not subtract out phonology and / or pragmatic and discourse processes identify the same network of regions (e.g., a contrast between natural spoken paragraphs and their acoustically degraded versions: Scott et al., 2016). Therefore, if we do not observe functional dissociations across language regions in their respective TRWs, it would not be simply because the language localizer subsamples a functionally homogeneous set of regions out of a larger network.

Second, to avoid over-inclusiveness, the contrast should not identify functional networks that are distinct from the core language network and might be recruited during online comprehension for other reasons (e.g., task-demands, attention, episodic encoding, non-verbal knowledge retrieval, or mental state inference). Whereas there are many potential differences between the processing of sentences *vs.* nonwords that might engage such non-linguistic processes, the identified regions exhibit robust language-selectivity in their responses, showing little or no response to non-linguistic tasks (Fedorenko et al., 2011; Fedorenko et al., 2012a; Pritchett et al., 2018; Ivanova et al., 2019; Jouravlev et al., 2019; For a review, see Fedorenko and Varley, 2016). Moreover, these regions are internally synchronized with one another during naturalistic cognition, yet are strongly dissociated from other brain networks (Blank et al., 2014; Paunov et al., 2019; for evidence from inter-individual differences, see: Mineroff et al., 2018). Therefore, if we observe functional dissociations across regions in their respective TRWs, it would not be simply because the localizer oversamples a functionally heterogeneous set of regions that extend beyond the core language network.

The evidence reviewed above provide strong support for both convergent and discriminant validity of the language localizer. In addition, this localizer generalizes across materials, tasks (passive reading *vs.* working memory), and modality of presentation (visual and auditory: Fedorenko et al., 2010; Braze et al., 2011; Vagharchakian et al., 2012). The latter is particularly important because the main task of the current study relied on spoken language comprehension.

#### 2.2.2. Critical experiment

Participants listened to the same materials that were originally used to characterize the cortical hierarchy of temporal integration windows. These materials were based on an audio recording of a narrated story (“Pie-Main”, told by Jim O’Grady at an event of “The Moth” group, NYC). They included (i) the intact audio; (ii) three “scrambled” versions of the story that differed in the temporal scale of incoherence, namely, lists of randomly ordered paragraphs, sentences, or words, respectively; and (iii) an audio-reversed version. The last stimulus served as a low-level control condition, because reverse speech contains the same acoustic characteristics as speech and is similarly processed (Lerner et al., 2011), but does not carry linguistic information beyond the phonetic level (Kimura and Folb, 1968; Koeda et al., 2006; but see Norman-Haignere et al., 2015).

To render these materials suitable for our existing scanning protocol, which used a repetition time of 2s (see section 2.3.1.), the silence and/or music period preceding and following each stimulus were each extended from 15s to 16s so that they fit an integer number of scans. These periods were not included in the analyses reported below. In addition, the two longest paragraphs in the paragraph-list stimulus were each split into two sections, and one section was randomly repositioned in the stream of shuffled paragraphs. No other edits were made to the original materials.

Participants listened to the materials over scanner-safe headphones (Sensimetrics, Malden, MA), in one of two orders: for 10 participants, the intact story was played first and was followed by increasingly finer levels of scrambling (from paragraphs to sentences to words). For the remaining 9 participants, the word-list stimulus was played first and was followed by decreasing levels of scrambling (from sentences to paragraphs to the intact story). The reverse story was positioned either in the middle of the scanning session or at the end, except for one participant for whom the scanning session had ended before we managed to play this stimulus.

At the end of the scanning session, participants answered 8 multiple-choice questions concerning characters, places and events from particular points in the narrative, with foils describing information presented elsewhere in the story. All participants demonstrated good comprehension of the story (17 of them answered all questions correctly, and the remaining two had only one error; the 20^th^ participant, excluded from analysis, had 50% accuracy).

### 2.3. Data acquisition and preprocessing

#### 2.3.1. Data acquisition

Structural and functional data were collected on a whole-body 3 Tesla Siemens Trio scanner with a 32-channel head coil at the Athinoula A. Martinos Imaging Center at the McGovern Institute for Brain Research at MIT. T1-weighted structural images were collected in 176 axial slices with 1mm isotropic voxels (repetition time (TR)=2530ms; echo time (TE)=3.48ms). Functional, blood oxygenation level-dependent (BOLD) data were acquired using an EPI sequence with a 90° flip angle and using GRAPPA with an acceleration factor of 2; the following parameters were used: thirty-one 4.4mm thick near-axial slices acquired in an interleaved order (with 10% distance factor), with an in-plane resolution of 2.1mm×2.1mm, FoV in the phase encoding (A>>P) direction 200mm and matrix size 96mm×96mm, TR=2000ms and TE=30ms. The first 10s of each run were excluded to allow for steady state magnetization.

#### 2.3.2. Data preprocessing

Spatial preprocessing was performed using SPM5 and custom MATLAB scripts. (Note that SPM was only used for preprocessing and basic first-level modeling, aspects that have not changed much in later versions; we used an older version of SPM because data for this study are used across other projects spanning many years and hundreds of participants, and we wanted to keep the SPM version the same across all the participants.) Anatomical data were normalized into a common space (Montreal Neurological Institute; MNI) template, resampled into 2mm isotropic voxels, and segmented into probabilistic maps of the gray matter, white matter (WM) and cerebrospinal fluid (CSF). Functional data were motion corrected, resampled into 2mm isotropic voxels, and high-pass filtered at 200s. Data from the localizer runs were additionally smoothed with a 4mm FWHM Gaussian filter, but data from the critical experiment runs were not, in order to avoid blurring together the functional profiles of nearby regions with distinct TRWs (we obtained the same results following spatial smoothing).

Additional temporal preprocessing of data from the critical experiment runs was performed using the CONN toolbox (Whitfield-Gabrieli and Nieto-Castanon, 2012) with default parameters, unless specified otherwise. Five temporal principal components of the BOLD signal time-courses extracted from the WM were regressed out of each voxel’s time-course; signal originating in the CSF was similarly regressed out. Six principal components of the six motion parameters estimated during offline motion correction were also regressed out, as well as their first time derivative. Next, the residual signal was bandpass filtered (0.008-0.09 Hz) to preserve only low-frequency signal fluctuations, because higher frequencies might be contaminated by fluctuations originating from non-neural sources (Cordes et al., 2001).

We note that bandpass filtering was not used by Lerner et al. (2011). When we repeated our analyses without including this step, we obtained the same pattern of results reported below. However, the unfiltered time-courses exhibited overall lower reliability across participants. We therefore chose to report the analyses of the filtered data.

### 2.4. Data analysis

All analyses were performed in Matlab (The MathWorks, Natick, MA) unless specified otherwise.

#### 2.4.1. Functionally defining language regions in individual participants

Data from the language localizer task were analyzed using a General Linear Model that estimated the voxel-wise effect size of each condition (sentences, nonwords) in each run of the task. These effects were each modeled with a boxcar function (representing entire blocks) convolved with the canonical Hemodynamic Response Function (HRF). The model also included first-order temporal derivatives of these effects, as well as nuisance regressors representing entire experimental runs and offline-estimated motion parameters. The obtained beta weights were then used to compute the voxel-wise *sentences > nonwords* contrast, and these contrasts were converted to *t*-values. The resulting *t*-maps were restricted to include only gray matter voxels, excluding voxels that were more likely to belong to either the WM or the CSF based on the probabilistic segmentation of the participant’s structural data.

Functional regions of interest (fROIs) in the language network were then defined using group-constrained, participant-specific localization (Fedorenko et al., 2010). For each participant, the *t*-map of the *sentences > nonwords* contrast was intersected with binary masks that constrained the participant-specific language regions to fall within areas where activations for this contrast are relatively likely across the population. These five masks, covering areas of the left temporal and frontal lobes, were derived from a group-level probabilistic representation of the localizer contrast in an independent set of 220 participants (available for download from: https://evlab.mit.edu/funcloc/download-parcels). In order to increase functional resolution in the temporal cortex, where a gradient of multiple different TRWs has originally been reported by Lerner et al. (2011), two temporal masks were each further divided in two, approximately along the posterior-anterior axis. The border locations marking these divisions were determined based on an earlier version of the group-level representation of the localizer contrast, obtained from on a smaller sample (Fedorenko et al., 2010). In total, seven masks were used (Figure 1), in the posterior, mid-posterior, mid-anterior, and anterior temporal cortex, the inferior frontal gyrus, its orbital part, and the middle frontal gyrus. (Unlike previous reports from our group that had used an additional mask in the angular gyrus, we decided to exclude this region going forward because it does not appear to be a part of the core language network in either its task-based responses or its signal fluctuations during naturalistic cognition. See, e.g., Blank et al., 2014; Blank et al., 2016; Pritchett et al., 2018; Ivanova et al., 2019; Jouravlev et al., 2019; Paunov et al., 2019).

**Figure 1.**
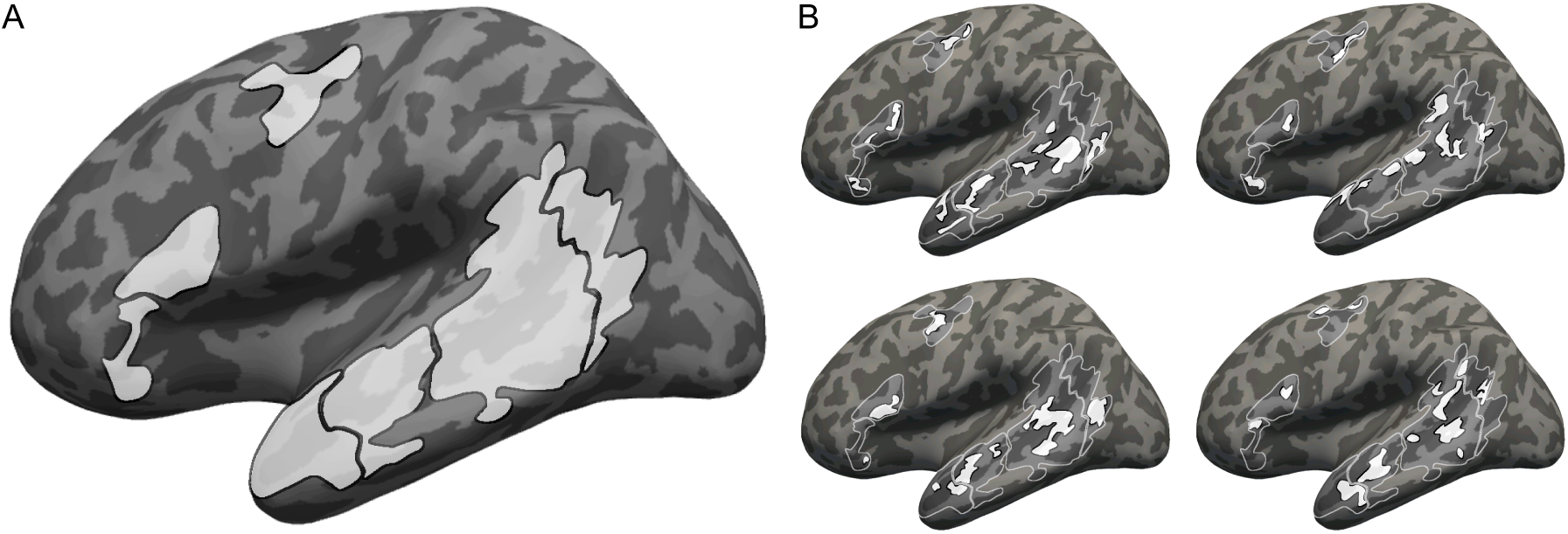
Defining participant-specific fROIs in the core language network. All images show approximate projections from functional volumes onto the surface of an inflated average brain in common (MNI) space. (A) Group-based masks used to constrain the location of fROIs. There masks were derived from a probabilistic group-level representation of the *sentences > nonwords* localizer contrast in a separate sample, following (Fedorenko et al., 2010). Contours of these masks are depicted in white in B. (B) Example fROIs of four participants. Note that, because data were analyzed in volume (not surface) form, some parts of a given fROI that appear discontinuous in the figure (e.g., separated by a sulcus) are contiguous in volumetric space.

In each of these masks, a participant-specific fROI was defined as the top 10% of voxels with the highest *t*-values for the *sentences > nonwords* contrast. fROIs within the smallest mask counted 37 voxels, and those within the largest mask—237 voxels (Figure 1). This top *n*% approach ensures that fROIs can be defined in every participant and that their sizes are the same across participants, allowing for generalizable results (Nieto-Castañón and Fedorenko, 2012).

We additionally defined a few alternative sets of fROIs for control analyses. First, to ensure that language fROIs were each functionally homogeneous and did not group together sub-regions with distinct TRWs, we also defined alternative, smaller fROIs based on the top 4% of voxels with the highest localizer contrast effects in each mask (these were 15-95 voxels in size). We were also interested in whether TRWs in the core language network differed from those in neighboring regions exhibiting weaker localizer contrast effects. Therefore, we defined fROIs based on the “second-best” 4% of voxels within each mask (i.e., those whose effect sizes were between the 92 and 96 percentile), as well as based on the “third best” (88-92 percentile), “fourth best” (84-88 percentile), and “fifth best” (80-84 percentile) sets.

#### 2.4.2. Main analysis of temporal receptive windows in language fROIs

For each of the five stimuli in the critical experiment (intact story, paragraph list, sentence list, word list, and reverse audio), in each of seven fROIs and for each participant, BOLD signal time-series were extracted from each voxel and were then averaged across voxels to obtain a single time-series per fROI, participant, and stimulus. When extracting these signals we skipped the first 6s (3 volumes) following stimulus onset, in order to exclude a potential initial rise of the hemodynamic response relative to fixation; such a rise would be a trivially reliable component of the BOLD signal that might blur differences across stimuli and fROIs. In addition, we included 6s of data following stimulus offset, in order to account for the hemodynamic lag.

To compute ISCs per fROI and stimulus, we *z-*scored the time-series of all participants but one, averaged them, and computed Pearson’s moment correlation coefficient between the resulting group-averaged time-series and the corresponding time-series of the left-out participant. This procedure was iterated over all partitions of the participant pool, producing 19 ISCs per fROI and stimulus. These ISCs were Fisher-transformed to improve the normality of their distribution (Silver and Dunlap, 1987).

To reiterate the logic detailed in the introduction, the resulting regional ISCs quantify the similarity of regional BOLD signal fluctuations across participants, with high values indicative of regional activity that reliably tracks the incoming input (correlations across participants mirror correlations within a single participant across stimulus presentations; Golland et al., 2007; Hasson et al., 2009; Blank and Fedorenko, 2017). Further, ISCs across the five stimuli constitute a functional profile characterizing a region’s TRW. Namely, reliable input tracking (i.e., high ISCs) is expected only for stimuli that are well-formed at the timescale over which a given region integrates information; weaker tracking (i.e., low ISCs) is expected for stimuli that are scrambled at that scale and, thus, cannot be reliably integrated.

For descriptive purposes, we first labeled the TRW of each fROI based on the most scrambled stimulus for which tracking was still statistically indistinguishable from tracking of the intact story. For example, if ISCs in a certain region were uniformly high for the intact story, paragraph list, and sentence list, but were significantly lower for the word list, that region’s TRW was labeled as “sentence-level” (because input tracking incurred a cost when well-formedness at that level was violated). Similarly, if ISCs in another region were uniformly high for the intact story, paragraph, sentence, and word lists, but dropped for the reverse audio, that region’s TRW was labeled as “word-level”. To thus label TRWs, for each fROI we compared ISCs between the intact story and every other stimulus using dependent-samples *t*-tests. The resulting *p*-values were corrected for multiple comparisons using false-discovery rate (FDR) correction (Benjamini and Yekutieli, 2001) across all pairwise comparisons and fROIs.

For our main analysis, we directly compared the pattern of ISCs to the five stimuli across the seven language fROIs via a two-way, repeated-measures analysis of variance (ANOVA) with fROI (7 levels) and stimulus (5 levels) as within-participant factors. The critical test was for a region-by-stimulus interaction. To further interpret our findings, we conducted follow-up analyses as detailed in the Results section. In addition to the parametric ANOVA, we ran empirical permutation tests of reduced residuals (Anderson and Braak, 2003), which are less sensitive to violations of the test’s assumptions, but obtained virtually identical results. We chose ANOVA over mixed-effects linear regression because the latter is more conservative due to estimator shrinkage, and we wanted to give any region-by-stimulus interaction—should one be present—the strongest chance of revealing itself. Nonetheless, inferences remained unchanged when tests were run via linear, mixed-effects regressions (using the lme4 toolbox in R) with varying intercepts by fROI, stimulus, and participant (Gelman, 2005).

#### 2.4.3. Controlling for baseline differences in input tracking across fROIs

When comparing functional responses across brain regions, it is critical to take into account regional differences in baseline responsiveness, because these might mask fROI-by-stimulus interactions (or expalin them away; Nieuwenhuis et al., 2011). For instance, whereas an ANOVA might conclude that a difference between an ISC of 0.5 for the intact story and an ISC of 0.4 for the reverse audio in one region is statistically indistinguishable from a difference between ISCs of 0.2 and 0.1 in another, the former difference constitutes only a 20% decrease whereas the latter constitutes a 50% decrease. Therefore, we corrected for such baseline differences and re-tested for a fROI-by-stimulus interaction.

To this end, we first regressed regional ISCs for the intact story out of ISCs for the other four stimuli in a linear, mixed-effects regression (using the lme4 toolbox in R). The model included varying intercepts and slopes by the following categorical variables: participant, fROI, stimulus, and the fROI-by-stimulus interaction. We used this full model so that variance associated with these variables (most importantly, with the interaction) would be correctly assigned to the corresponding varying intercepts rather than be mistakenly attributed to baseline input tracking (i.e., the fixed effect of ISCs for intact story, or its associated varying slopes). Next, we extracted from the model estimates of ISCs in each region for the paragraph list, sentence list, word list, and reverse audio stimuli, but these estimates were based only on coefficients for the fixed effect of the intact story and its varying slopes by the different variables. Coefficients for varying intercepts by each categorical variable were ignored, to avoid subtracting the effects of these variables out of the data. Finally, the extracted estimates were subtracted from their respective veridical ISCs to obtain residuals that were not contaminated by regional differences in baseline input tracking. We tested these residuals for fROI-by-stimulus interaction, as described in section 2.4.2.

#### 2.4.4. Additional, voxel-based analyses

Whereas our main analyses examined ISCs in functionally defined, participant-specific fROIs, we also conducted two control analyses for which ISCs were defined using the common, voxel-based approach. Although we believe this approach is disadvantageous and suffers from interpretational limitations (see Introduction), we chose to use it nonetheless to provide a more comprehensive investigation of TRWs in the core language network.

Our first goal was to replicate the original findings by Lerner et al. (2011) so as to ensure that any differences between our main analyses and this previous study were not due to inconsistencies in the data. For this analysis, in line with Lerner et al. (2011), we smoothed the (temporally preprocessed) functional scans with a 6mm FWHM Gaussian kernel. Then, for each of the five stimuli in the main experiment, we computed voxel-wise ISCs for the subset of left-hemispheric coordinates that met the following three criteria: (i) were more likely to be gray matter than either WM or CSF in at least 2/3 of the participants, based on the probabilistic segmentation of their individual, structural data; (ii) were part of the frontal, temporal, or parietal lobes as defined by the AAL2 atlas (Tzourio-Mazoyer et al., 2002; Rolls et al., 2015); and (iii) fell in the cortical mask used for the cortical parcellation in Yeo et al. (2011). We then labeled the TRW of each voxel following the approach of Lerner et al. (2011), namely, based on the most scrambled stimulus that, across participants, was still tracked significantly above chance (evaluated against a Gaussian fit to an empirical null distribution of ISCs that was generated from surrogate signal time-series; see Theiler et al., 1992). Tests were FDR corrected for multiple comparisons across stimuli and voxels. The resulting map of TRWs was projected onto Freesurfer’s average cortical surface in MNI space. No further quantitative tests were performed, as we were only interested in obtaining a visually similar gradient of TRWs to the one previously reported.

Our second goal was an alternative definition of TRWs that relied less heavily on the task-based functional localizer, in order to alleviate any remaining concerns regarding its use (see section 2.2.1.). Here, rather than defining fROIs that maximized the localizer contrast effect and subsequently characterizing their TRWs, we aimed at defining fROIs that directly maximized the ISC profiles consistent with certain TRWs. To avoid circularity from using the same data to define fROIs and to estimate their response profiles (Vul and Kanwisher, 2010; Kriegeskorte et al., 2009), we first created two independent sets of ISCs by splitting each BOLD signal time-series and computing ISCs for each half of the data. We then used data from the second half of each stimulus to define fROIs, and data from the first half to compare TRWs across the resulting fROIs (We used the first half of each stimulus for the critical test because we suspected input tracking would be overall lower in the second half due to participants losing focus, especially for scrambled stimuli without coherent meaning; however, the pattern of results did not depend on which data half was used for fROI definition and which was used for the critical test.)

This analysis proceeded as follows: in each mask, we labeled the TRW of each voxel as in our main analysis (section 2.4.2.), i.e., based on the most scrambled stimulus for which tracking was statistically indistinguishable from tracking of the intact story (the alternative labeling scheme described in the previous paragraph, based on the most scrambled stimulus that was still tracked significantly above chance, yielded the same results). Next, we focused on voxels whose TRW label was the same as the label of the localizer-based fROI from the main analysis (focusing, instead, on the TRW label whose voxels showed the strongest responsiveness to sentence reading compared to fixation in the localizer task yielded the same results). We sorted these voxels based on the difference between their tracking of the intact story and of the least scrambled stimulus that was tracked less reliably, and chose the 27 voxels whose *p*-values for that comparison were the smallest (most significant). For example, if the TRW of interest was “sentence-level”, this meant that we focused on voxels (i) whose tracking of the sentence list did not statistically differ from their tracking of the intact story, but (ii) their tracking of the word list was significantly less reliable; a “sentence-level” fROI was then chosen as the 27 voxels showing the most significant differences between ISCs for the intact story and the word list stimulus (in a dependent-samples *t*-test across participants). Once a fROI was defined this way in each mask, we averaged the ISCs for the first half of each stimulus across its voxels, and compared the resulting ISC profiles across fROIs using ANOVA tests as in our main analyses (section 2.4.2.).

#### 2.4.5. Comparing TRWs between the language network and other functional regions

To situate data from the core language network in a broader context, we also computed ISCs in other cortical regions. First, we computed ISCs based on BOLD signal time-series averaged across voxels from an anatomically defined mask of lower-level auditory cortex in the anterolateral section of Heschl’s gyrus in the left hemisphere (Tzourio-Mazoyer et al., 2002) (we did not use a functional localizer because lower-level sensory regions show overall better mapping onto macro-anatomy compared to higher-level associative regions). This region has a fast temporal timescale (e.g., Lerner et al., 2011; Honey et al., 2012a), and was expected to track all stimuli equally reliably.

Second, we extracted ISCs from regions of the “episodic” (or “default mode”) network (Gusnard and Raichle, 2001; Raichle et al., 2001; Buckner et al., 2008; Andrews-Hanna et al., 2010; Humphreys et al., 2015), which is engaged in processing episodic information. This network, recruited during narrative construction, is expected to integrate input over longer timescales compared to the core language network (Regev et al., 2013; Chen et al., 2016; Margulies et al., 2016; Simony et al., 2016; Yeshurun et al., 2017a; Yeshurun et al., 2017b; Zadbood et al., 2017; Nguyen et al., 2019). To identify fROIs in this network we relied on its profile of deactivation during tasks that tax executive functions for the processing of external stimuli, and used a visuo-spatial working-memory localizer that included a “hard” condition requiring the memorization of 8 locations on a 3×4 grid (Fedorenko et al., 2011; Fedorenko et al., 2013). We defined the contrast *hard < fixation* and, following the approach outlined above (section 2.4.1.), chose the top 10% of voxels showing the strongest *t*-values for this contrast in two left-hemispheric masks, located in the posterior cingulate cortex and temporo-parietal junction (these masks were generated based on a group-level probabilistic representation of the localizer task data from 197 participants).

To compare TRWs between the core language network and each of these two other systems, we averaged ISCs for each stimulus across fROIs in each system and conducted two-way, repeated-measures ANOVAs with system (2 levels: language and auditory / language and episodic) and stimulus (5 levels) as within-participant factors. The critical test was for a system-by-stimulus interaction.

## 3. Results

### 3.1. No evidence for a region-by-stimulus interaction across temporal and inferior frontal language fROIs

#### 3.1.1. Main analysis

The main results of the current study are presented in Figure 2A. We computed inter-subject correlations for each of the five stimuli in each of seven core language fROIs, and directly compared the resulting regional profiles of input tracking via a two-way (fROI×stimulus), repeated-measures ANOVA. As expected, there was a main effect of stimulus (*F*_(68,4)_=32.7, partial *η*^2^=0.66, *p*<10^−14^). Follow-up ANOVAs (FDR corrected for multiple comparisons) contrasting the intact story to each other stimulus revealed no overall differences in tracking the intact story and paragraph list (*F*_(1,18)_=0, *η*^2^_*p*_=0, *p*=1) or sentence list (*F*_(1,18)_=0.02, *η*^2^_*p*_=10^−3^, *p*=1), but weaker tracking of the word list (*F*_(1,18)_=27.27, *η*^2^_*p*_=0.6, *p*<10^−3^) and reverse audio (*F*_(1,18)_=169.90, *η*^2^_*p*_=0.92, *p*<10^−7^). Furthermore, the sentence list was tracked more reliably than the word list (*F*_(1,18)_=9.76, *η*^2^_*p*_=0.35, *p*=0.04) which was, in turn, tracked more reliably than the reverse audio (*F*_(1,18)_=48.56, *η*^2^_*p*_=0.74, *p*<10^−4^). In addition, there was a main effect of fROI (*F*_(6,102)_=19.7, *η*^2^_*p*_=0.54, *p*<10^−14^), indicating that some fROIs overall tracked stimuli more strongly than others, a finding we return to in section 3.1.2.

**Figure 2.**
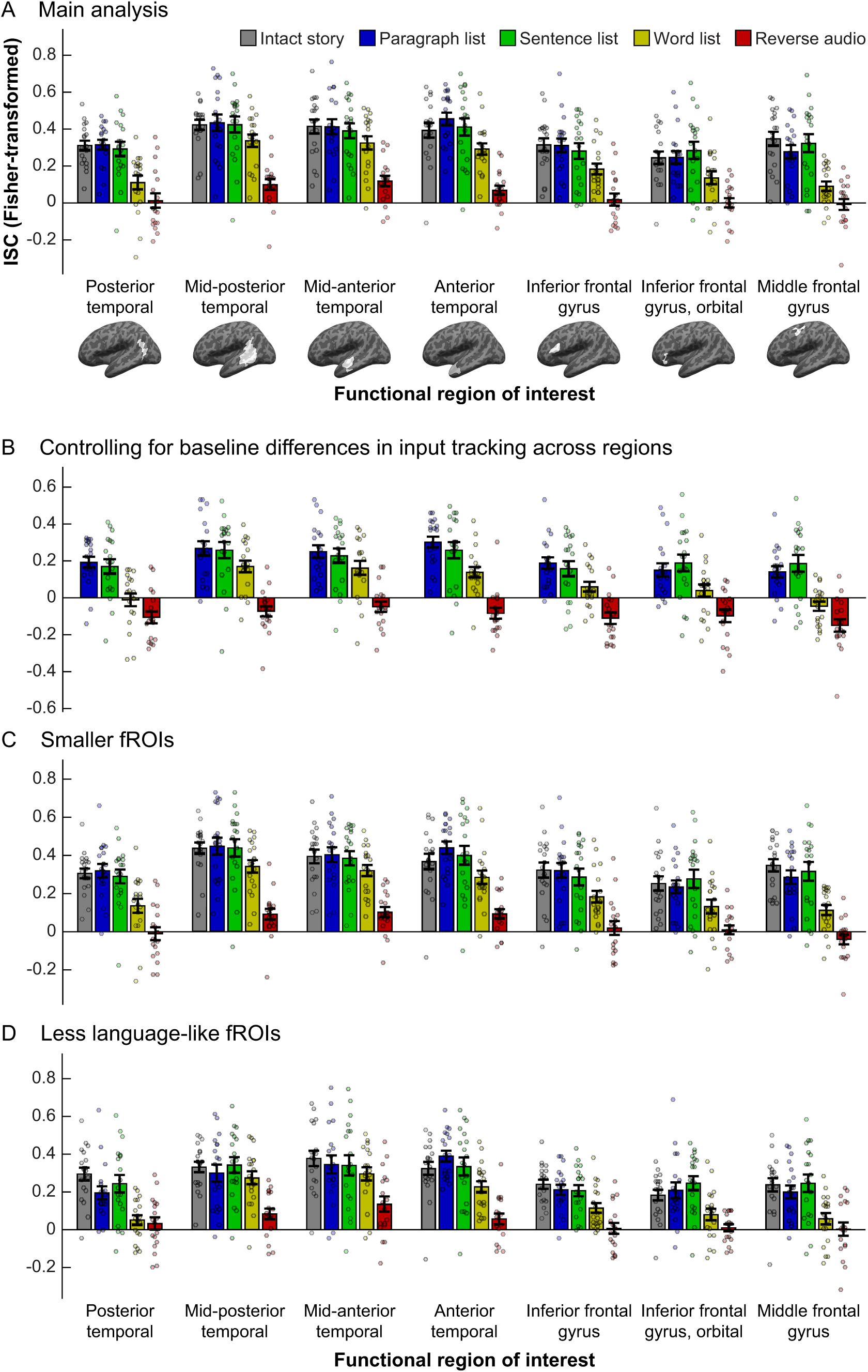
Comparing temporal receptive windows (TRWs) across core language regions. In each panel, inter-subject correlations (ISCs, a proxy for input tracking reliability; *y*-axis) are shown for each stimulus (bar colors) in each fROI (*x*-axis; brain images depict whole masks, but data are for participant-specific fROIs defined within those masks). Dots show individual data points, bars show means across the sample, and error-bars show standard errors of the mean. (A) Main analysis. (B) Residualized ISCs after regressing out the ISC of the intact story (a “baseline” measure of tracking). (C) ISCs in smaller fROIs, defined as the top 4% of voxels in each mask that showed the biggest *sentences > nonwords* effect in the localizer task, rather than the top 10% as in (A). (D) fROIs defined as the “fifth-best” 4% of voxels in each mask, i.e., those whose effect size for the localizer contrast was in the 80-84 percentile range.

As an initial characterization of the region-wise TRWs, we ran dependent-samples *t*-tests in each fROI to compare ISCs for the intact story and for each of the other stimuli. When then identified the most scrambled stimulus whose tracking was still statistically indistinguishable from tracking of the intact story (correcting for multiple tests across pairwise comparisons and fROIs). These tests indicated that the mid-posterior, mid-anterior, and anterior temporal fROIs each exhibited “word-level” TRWs, with input tracking reliability not incurring a cost when words were randomly ordered, but becoming significantly weaker for the reverse audio (for all three regions: *t*_(17)_>6.54, Cohen’s *d*>1.58, *p*<10^−4^). The posterior temporal cortex, inferior frontal gyrus, its orbital part, and middle frontal gyrus each exhibited “sentence-level” TRWs, with input tracking reliability not incurring a cost when sentences were randomly ordered, but becoming significantly weaker when word order within sentences was scrambled (for all four regions: *t*_(18)_>2.8, *d*>0.66, *p*<0.04). Prior to multiple comparison correction, all seven fROIs exhibited “word-level” TRWs. Based on these findings, in several of our analyses below we report tests focusing on the sentence list and word list conditions, which appear to be the locus of potential functional differences across fROIs.

Critically, whereas the ANOVA test for a fROI-by-stimulus interaction in ISCs was significant (*F*_(24,408)_=1.78, *η*^2^_*p*_=0.10, *p*=0.014), this interaction was explained by the middle frontal gyrus (MFG): follow-up analyses testing for an interaction across all regions but one (uncorrected for multiple comparisons, so as to be anti-conservative) failed to reach significance when the MFG was removed (*F*_(20,340)_=1.32, *η*^2^_*p*_=0.07, *p*=0.16), but remained significant when each of the other fROIs was removed (for all tests, *F*_(20,340)_>1.71, *η*^2^_*p*_>0.09, *p*<0.03). The same results obtained when testing only the sentence list and word list stimuli (all seven fROIs: *F*_(6,108)_=4.32, *η*^2^_*p*_=0.19, *p*<10^−3^; MFG excluded: *F*_(5,90)_=2.24, partial *η*^2^=0.11, *p*=0.057; any other fROI excluded: *F*_(5,90)_>4.11, *η*^2^_*p*_>0.18, *p*<0.002). Indeed, the word list stimulus was driving the interaction across the seven fROIs: when this stimulus was excluded from analysis, the interaction test failed to reach significance (*F*_(18,306)_=1.13, *η*^2^_*p*_=0.06, *p*=0.32), but it did remain significant when each of the other stimuli were removed (reverse audio excluded: *F*_(18,324)_=2.56, *η*^2^_*p*_=0.13, *p*<10^−3^; any other stimulus excluded: *F*_(18,306)_>1.82, *η*^2^_*p*_=0.10, *p*<0.03).

Below, we report several additional analyses exploring the fROI-by-stimulus interaction (or lack thereof). Taken together, these analyses find no evidence that temporal and inferior frontal fROIs have functionally distinct temporal receptive windows.

#### 3.1.2. Controlling for baseline differences in input tracking across fROIs

Language fROIs differed from one another in their “baseline” input tracking: a one-way ANOVA performed on the intact-story ISCs with fROI (7 levels) as a within-participant factor revealed a significant main effect (*F*_(6,108)_=5.13, *η*^2^_*p*_=0.22, *p*=10^−4^). To control for these differences, we residualized out the ISCs of the intact story from ISCs of the other four stimuli, and re-ran the ANOVA testing for fROI-by-stimulus interaction on the residuals of the latter stimuli (Figure 2B). As in our main analysis, this test revealed a fROI-by-stimulus interaction (*F*_(18,306)_=1.83, *η*^2^_*p*_=0.10, *p*=0.02) that was accounted for by the MFG (MFG excluded: *F*_(15,255)_=1.51, *η*^2^_*p*_=0.08, *p*=0.10). The results obtained when testing only the sentence list and word list stimuli were virtually identical to those of the corresponding test reported above (all seven fROIs: *F*_(6,108)_=4.32, *η*^2^_*p*_=0.19, *p*<10^−3^; MFG excluded: *F*_(5,90)_=2.24, *η*^2^_*p*_=0.11, *p*=0.057). Therefore, it is unlikely that evidence for distinct TRWs across language fROIs was “washed out” by regional differences in baseline input tracking.

#### 3.1.3. Testing smaller fROIs

The same results as in the main analysis were obtained when we tested smaller fROIs defined as the top 4% (rather than 10%) of voxels showing the strongest localizer contrast effects in each mask (Figure 2C): when all 7 fROIs and 5 stimuli were included, there was a fROI-by-stimulus interaction (*F*_(24,408)_=1.62, *η*^2^_*p*_=0.09, *p*=0.03), which was no longer significant once the MFG was excluded (*F*_(20,340)_=1.20, *η*^2^_*p*_=0.07, *p*=0.25). Similarly, when we tested only the sentence list and word list stimuli, there was a significant fROI-by-stimulus interaction across the 7 fROIs (*F*_(6,108)_=2.80, *η*^2^_*p*_=0.13, *p*<0.02) but not across the six fROIs excluding the MFG (*F*_(20,340)_=1.48, *η*^2^_*p*_=0.08, *p*=0.21). Thus, the lack of evidence for a fROI-by-stimulus interaction in our main analysis is unlikely to result from using regions that were too large and grouped together several, functionally distinct sub-regions.

#### 3.1.4. Testing less language-like fROIs

The lack of evidence for a fROI-by-stimulus interaction across core language regions might have resulted from lack of power to detect such interactions. To examine this possibility, we conducted the same analysis on each of several sets of alternative fROIs that, instead of showing the strongest *sentences > nonwords* effects, consisted of voxels that showed weaker localizer contrast effects. Specifically, the localizer contrast effects in these voxels were either in the 92-96 percentiles of their respective masks (“second-best” 4%), 88-92 percentiles, 84-92 percentiles, or 80-84 percentiles (“third-”, “fourth-”, and “fifth-best” 4%, respectively). We reasoned that such voxels, which showed less language-like responses, were either more peripheral members of the language network, or belonged to other functional networks that lie in close proximity to language regions (Chein et al., 2002; Fedorenko et al., 2012a; Deen et al., 2015). As such, these regions might differ from one another, and from the core language fROIs, in their integration timescales.

In each of these alternative sets of 7 fROIs, a two-way, repeated-measures ANOVA revealed a fROI-by-stimulus interaction, indicating that these regions differed from one another in their TRWs (second-best set: *F*_(24,408)_=1.87, *η*^2^_*p*_=0.09, *p*=0.007; third-best set: *F*_(24,408)_=1.95, *η*^2^_*p*_=0.10, *p*=0.005; fourth-best set: *F*_(24,408)_=1.92, *η*^2^_*p*_=0.10, *p*=0.006; fifth-best set: *F*_(24,408)_=1.95, *η*^2^_*p*_=0.10, *p*=0.006). For the fROIs consisting of second-best voxels—voxels that, being within the top 10%, were part of the fROIs used for the main analysis—this interaction was driven by the MFG (fROI-by-stimulus interaction with MFG excluded: *F*_(20,340)_=1.50, *η*^2^_*p*_=0.08, *p*=0.08). In contrast, for the other sets of fROIs, the interaction remained significant even with the MFG removed, and its effect size descriptively grew as less language-like fROIs were tested (fROI-by-stimulus interaction with MFG excluded, third-best set: *F*_(20,340)_=1.78, *η*^2^_*p*_=0.09, *p*=0.02; fourth-best set: *F*_(20,340)_=1.96, *η*^2^_*p*_=0.10, *p*=0.008; fifth-best set: *F*_(20,340)_=2.35, *η*^2^_*p*_=0.12, *p*=10^−3^) (Figure 2D). These findings suggest that our study had enough power to detect fROI-by-condition interactions when those exist; such interaction is simply not evident in the core language network, whose regions show indistinguishable TRWs.

### 3.2. Additional, voxel-based analyses support the main finding

To demonstrate that the lack of evidence for a fROI-by-stimulus interaction in the core language network was not constructed into the findings via our choice of localizer task, we re-computed ISCs using the common, voxel-by-voxel approach. First, we observed that the resulting ISCs qualitatively replicate the overall topography TRWs reported by Lerner et al. (2011) (Figure 3A). Namely, large portions of the superior temporal cortex exhibit reliable tracking of all five stimuli, including the reverse audio, indicative of a short TRW; middle temporal regions reliably track word lists (as well as less scrambled stimuli) but not the reverse audio, i.e., are sensitive to well-formedness on the timescale of morphemes or words; extending further in inferior, posterior, and anterior directions, some temporal and parietal regions exhibit longer TRWs and reliably track only stimuli whose structure is well-formed at the level of phrases or sentences; and in the frontal lobe, voxels exhibit sensitivity to coherence at either the word-, sentence-, or paragraph-level. The broad consistency between this pattern and the previously established hierarchy of integration timescales indicates that the lack of functional dissociations across core language regions cannot be attributed to fundamental inconsistencies between the current data and those of Lerner et al. (2011).

**Figure 3.**
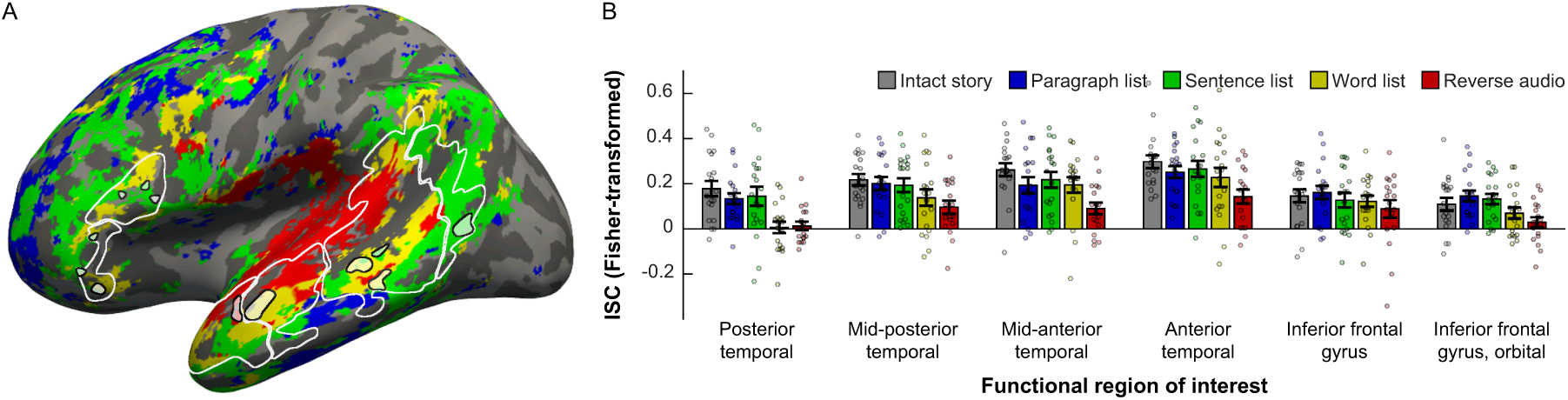
Voxel-based analyses. (A) Voxel-based ISCs in the frontal, parietal, and temporal lobes of the left-hemisphere are projected from a functional volume onto the surface of an inflated average brain in common (MNI) space. Each voxel is colored according to its TRW, defined as the most scrambled stimulus that, across participants, was still tracked significantly above chance (following the method in Lerner et al., 2011) (see legend in (B)). Note the progression from a short TRW (significant tracking of all stimuli, labeled in red), around the mid-posterior and mid-anterior temporal lobe, towards longer TRWs (yellow or green) in surrounding temporal areas, and finally to the frontal lobe (yellow, green, or blue). White contours mark the masks that were used for the main analysis (see Figure 1), excluding the MFG. Small black contours surrounding faint colors mark a set of fROIs that was defined as an alternative to the main fROIs, by choosing the top 27 voxels in each mask showing the strongest pattern consistent with that mask’s TRWs as identified in the main analysis (see Figure 2A). These voxels were chosen based on contrasting ISCs for the intact story to ISCs for each of the other stimuli, and relied on data from only the second half of each stimulus (unlike the coloring scheme in this panel, which is based on data from entire time-series of each stimulus and compared the ISCs of each stimulus against chance). (B) Cross-validated TRWs of the fROIs marked in (A), based on ISCs for data from only the first half of each stimulus. Conventions are the same as in Figure 2.

Next, as a final attempt at uncovering distinct integration timescales across temporal and inferior frontal language regions, we defined an alternative set of fROIs by directly searching for certain TRWs. Recall that, in our descriptive labeling of fROIs in the main analysis (section 3.1.1), two functional profiles were observed: on the one hand, fROIs in the mid-posterior, mid-anterior, and anterior temporal areas exhibited sensitivity to morpheme/word-level information, with input tracking incurring a significant cost only for the reverse audio. On the other hand, fROIs in the posterior temporal, inferior frontal, and orbital areas exhibited sensitivity to phrase/sentence-level information, with input tracking incurring a cost not only for the reverse audio but also for the word list. Now, we aimed at maximizing the differences between these two profiles, which did not reliably differ from one another in the main analysis. For this purpose, we defined the following fROIs: (i) within each of the former three masks, among those voxels whose ISCs did not significantly differ between the intact story and word list, we selected the 27 voxels with the biggest difference between ISCs for the intact story and the reverse audio; (ii) within each of the latter three masks, among those voxels whose ISCs did not significantly differ between the intact story and sentence list, we selected the 27 voxels with the biggest difference between ISCs for the intact story and the word list.

These six fROIs were defined based on data from the second half of each stimulus, and we then conducted a two-way, repeated-measures ANOVA to test for a fROI-by-stimulus interaction in ISCs from the first half of each stimulus (Figure 3B). The interaction was not significant (*F*_(20,340)_=1.26, *η*^2^_*p*_=0.07, *p*=0.21). When testing only the sentence list and word list stimuli, the interaction was very weak (*F*_(5,90)_=2.35, *η*^2^_*p*_=0.12, *p*=0.047), especially considering that the choice of which TRW to define in each mask (based on the main analysis) relied on the same functional data tested here (even though, within the current test itself, fROI definition and response estimation were performed on two independent halves of those data). Similar results were obtained when fROIs were defined based on data from the first half of each stimulus and the ANOVA was run on ISCs from the second half (sentence list and word list only: *F*_(5,90)_=1.54, *η*^2^_*p*_=0.08, *p*=0.19).

### 3.3. The core language network as a unified whole occupies a unique step within a broader cortical hierarchy of integration timescales

The finding that core language regions in temporal and inferior frontal areas show indistinguishable TRWs does not challenge the hypothesis of a broader hierarchy of integration timescales throughout the cortex. Rather, it indicates that core language regions do not occupy multiple, distinct stages within this hierarchy. Yet other functional regions occupy other stages, some with shorter TRWs than those of the language network and others with longer TRWs.

To demonstrate this, we first compared the ISCs for the five stimuli averaged across six language fROIs (excluding the MFG) to ISCs from the auditory cortex. A two-way, repeated-measures ANOVA yielded a system (language *vs.* auditory) by stimulus interaction (*F*_(4,68)_=10.53, *η*^2^_*p*_=0.38, *p*<10^−5^) (Figure 4). We followed up on this result with system-by-stimulus interaction tests that only included the intact story and one other stimulus. These tests (FDR corrected for multiple comparisons) revealed that, compared to the core language network, ISCs in the auditory region differed less between the intact story and the word list (*F*_(1,18)_=9.75, *η*^2^_*p*_=0.35, *p*=0.025), as well as between the former stimulus and the reverse audio (*F*_(1,17)_=34.11, *η*^2^_*p*_=0.67, *p*<10^−3^). In other words, input tracking in the auditory region incurred lower costs for fine-grained scrambling, indicative of a shorter TRW compared to that of the core language network.

**Figure 4.**
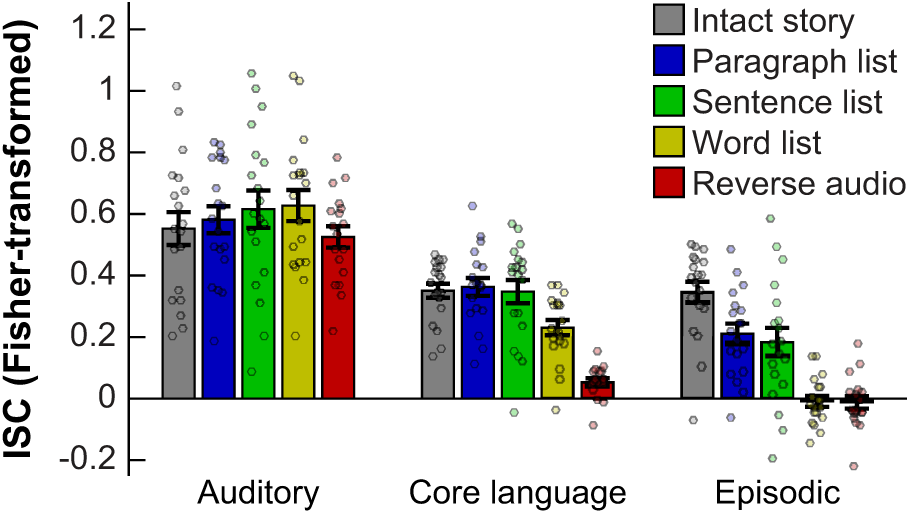
Demonstrating a hierarchy of TRWs across functionally distinct brain regions. ISCs are shown for a low-level auditory region (around the antero-lateral portion of Heschl’s gyrus; left), the core language network (averaged across fROIs, excluding the MFG; middle), and a subset of the episodic network (averaged across fROIs in the left posterior cingulate cortex and temporo-parietal junction; right). Conventions are the same as in Figure 2.

Next, we similarly compared ISCs in the language network to those in the episodic network (averaged across left-hemisphreric posterior cingulate and temporo-parietal fROIs). Again, we found a network by stimulus interaction (*F*_(4,68)_=10.82, *η*^2^_*p*_=0.69, *p*<10^−6^) (Figure 4). Follow up interaction tests revealed that, compared to the core language network, ISCs in the episodic network differed more between the intact story and the paragraph list (*F*_(1,18)_=21.30, *η*^2^_*p*_=0.54, *p*=10^−3^), sentence list (*F*_(1,18)_=15.95, *η*^2^_*p*_=0.47, *p*=0.005), and word list (*F*_(1,18)_=43.68, *η*^2^_*p*_=0.71, *p*<10^−4^). Input tracking in the episodic network thus incurred higher costs for coarse violations of well-formedness on the timescale of paragraph and sentences and, further, showed no tracking of word lists. This functional profile is indicative of a longer TRW compared to that of the core language network.

These findings situate the core language network within the context of a cortical hierarchy of integration timescales (Himberger et al., 2018). The common functional profile shared by temporal and inferior frontal language regions occupies a particular stage within this broader hierarchy, which is located, as expected, downstream from auditory regions and upstream from the episodic network.

## 4. Discussion

The current study examined how reliably different regions in the core language network track linguistic inputs that violated well-formedness at various levels. To this end, we recorded regional time-series of BOLD signal fluctuations elicited by increasingly scrambled versions of a narrated story, and measured the reliability of these fluctuations across individuals to quantify the extent to which they were stimulus-locked. We found that patterns of input tracking in left temporal and inferior frontal language regions all exhibited statistically indistinguishable profiles of sensitivity to linguistic structure at different grain sizes. Namely, these regions all tracked paragraph lists and sentence lists as reliably as they tracked the intact story, but tracked word lists less reliably, and tracked a reverse audio only weakly or not at all. These findings suggest that language regions integrate information over a common timescale, window, which is (i) sensitive to structure at or below the word level, given the increased tracking of the word list compared to the reverse audio; (ii) also sensitive to structure at the phrase or sentence level, given the further increase in tracking of the sentence list compared to the word list; but (iii) not sensitive to information above the sentence level, given no further boosts in tracking of the paragraph list compared to the sentence list. This common profile of information integration provides a novel functional signature of perisylvian, high-level language regions.

We emphasize that our main, null findings of a region-by-stimulus interaction constitute lack of evidence, and not evidence for a lack of functional dissociation across the core language network. Nevertheless, we extensively tested and rejected alternative explanations for these null results: they are not likely to be accounted for by baseline differences in input tracking across regions, which could have masked differences in TRWs (section 3.1.2.); by fROIs being large enough to include—and average across—multiple, functionally distinct sub-regions (section 3.1.3.); by lack of power to detect region-by-stimulus interactions in the general cortical areas we focused on (section 3.1.4.); or by relying on a task-based, functional localizer to identify participant-specific regions of interest (section 3.2.). We therefore conclude that no compelling evidence have been found in favor of a functional dissociation across core language regions in terms of their integration timescales.

The current results are inconsistent with a division of linguistic labor across the core language network that is topographically organized by integration timescales (cf. Lerner et al., 2011; DeWitt and Rauschecker, 2012; Bornkessel-Schlesewsky et al., 2015; Hasson et al., 2015; Chen et al., 2016; Baldassano et al., 2017; Yeshurun et al., 2017a; Sheng et al., 2018). Instead, they support the hypothesis that temporal and inferior frontal core language regions form a unified whole that occupies a unique stage within a broader cortical hierarchy of temporal integration. In this cortical hierarchy, core language regions follow lower-level speech-processing regions (Mesgarani et al., 2014; Overath et al., 2015; Poeppel, 2003; Vagharchakian et al., 2012), and precede higher-level associative regions that integrate full narratives (Regev et al., 2013; Chen et al., 2016; Margulies et al., 2016; Simony et al., 2016; Yeshurun et al., 2017a; Yeshurun et al., 2017b; Zadbood et al., 2017; Nguyen et al., 2019). The only region of the language network that might occupy a different stage in this hierarchy is the middle frontal gyrus, which has a somewhat longer integration timescale compared to the rest of the core language network.

### 4.1. Evidence for a distributed cognitive architecture of language processing

The finding of a common integration window shared across temporal and inferior frontal core language regions constrains comprehension models, challenging the notion of a functional dissociation between processes that either operate on different timescales and/or construct linguistic representations of different grain sizes. Instead, it suggests that linguistic processes on multiple timescales—from morpheme- or word-level to phrase- or sentence-level—are implemented in neural circuits that are distributed rather than focal and, moreover, overlap with one another and are thus cognitively inseparable. This view is consistent with psycholinguistic theories (Joshi et al., 1975; Schabes et al., 1988; Goldberg, 1995; Bybee, 1998; Jackendoff, 2002; Culicover and Jackendoff, 2005; Wray, 2005; Bybee, 2010; Snider and Arnon, 2012; Jackendoff, 2007; Langacker, 2008; Christiansen and Arnon, 2017) and empirical evidence (Clifton et al., 1984; MacDonald et al., 1994; Trueswell et al., 1994; Garnsey et al., 1997; Traxler et al., 2002; Farmer et al., 2006; Reali and Christiansen, 2007; Bradlow and Bent, 2008; Gennari and MacDonald, 2008; Maye et al., 2008; Trude and Brown-Schmidt, 2012; Schmidtke et al., 2014) in favor of a representational continuum that does away with traditionally posited boundaries and extends from phonemes, to morphemes, to words with their syntactic and semantic attributes, to phrase constructions and their meanings. A continuous gradient rather than a strict hierarchy of distinct stages is supported by the observation that different language regions exhibited TRWs that, while not statistically distinguishable, were nonetheless somewhat descriptively different from one another.

The current data do not exclude the possibility that temporal and inferior frontal core language regions, which all show the same functional profile in terms of integration timescale, each implement a distinct set of processes relevant to such integration (see, e.g., Hagoort and Indefrey, 2014; Brennan et al., 2016). However, our finding of a functional signature distributed across the language network adds to prior neuroimaging studies reporting overlapping and distributed activations across diverse linguistic tasks (Gernsbacher and Kaschak, 2003; Démonet et al., 2005; Vigneau et al., 2006; Price, 2012). These include tasks of phonological (Scott and Wise, 2004; Hickok and Poeppel, 2007; Turkeltaub and Coslett, 2010), lexical (Paulesu et al., 1993; Indefrey and Levelt, 2004; Blumstein, 2009; Anderson et al., 2018), syntactic (Caplan, 2007; Bautista and Wilson, 2016; Blank et al., 2016), and semantic (Bookheimer, 2002; Thompson-Schill, 2003; Patterson et al., 2007; Binder et al., 2009; Fedorenko et al., 2016; Fedorenko et al., 2018; Mollica et al., 2018; Siegelman et al., 2019) processing. Unlike these traditional, task-based studies, which used cleverly controlled manipulations contrived to isolate particular aspects of linguistic processing, the current study employed an alternative approach (Hasson et al., 2004) based on richly structured stimuli in a naturalistic listening paradigm. It therefore importantly complements the prior evidence for a distributed architecture for language processing within which the very same neural circuits support many different kinds of computations.

### 4.2. Why functional dissociations among language regions might go undetected

While we interpret our findings as supporting the distributed implementation of language processing in overlapping neural circuits, they are not inconsistent with some forms of functional dissociations across distinct linguistic mechanisms. Indeed, neuropsychological findings, despite their many inconsistencies, indicate that at least some language regions are likely specialized for some linguistic processes and not others, because some people with aphasia following brain lesions drastically differ from one another in their behavioral symptoms (Caramazza and Coltheart, 2006). It is thus possible that some fMRI evidence for distributed linguistic processing underestimate a more complex functional architecture within the language network.

Several forms of functional dissociations could have gone undetected in the current study. First, the relatively uncontrolled properties of the naturalistic paradigms and the substantial differences across stimuli in both the localizer and main tasks might have been unsuitable for detecting subtler linguistic distinctions; and the definition of a single participant-specific fROI in each mask might have compromised our ability to identify distinctions among small regions that lie in close proximity to one another (e.g., Humphries et al., 2005; Hagoort, 2014; Wilson et al., 2018). Nonetheless, whether such distinctions as previously reported in the literature are replicable, and whether they reflect language-specific functions rather than more general, conceptual processes, remains debated (For one such example, see Dapretto and Bookheimer, 1999; Siegelman et al., 2019; for another, see Frankland and Greene, 2015; Wang et al., 2016; Anderson et al., 2018; see also Vigliocco et al., 2011; Moseley and Pulvermüller, 2014).

Second, due to spatial resolution limits of fMRI, dissociations at the sub-voxel level could be missed, i.e., functional profiles of distinct neural circuits that are all located within a single voxel would be aggregated together. Decomposing voxel-level BOLD signals into distinct components would require more sophisticated analytical techniques than the ones used here (e.g., Norman-Haignere et al., 2015). Nevertheless, given that the neuropsychological literature has studied many patients with lesions much larger than the spatial grain of fMRI, at least some functional subdivisions within the language network should in principle be detectable across regions rather then within voxels.

Third, the low temporal resolution of the fMRI BOLD signal limits the ability to detect functional distinctions in the time or frequency domains. The neural tracking of linguistic information rapidly evolves over just hundreds of milliseconds (Gross et al., 2013; Ding et al., 2016), but the hemodynamic response to neural activity reaches a peak only after several seconds, smoothing over any putative differences in the timing at which distinct linguistic processes could engage a given region. Similar concerns apply to distinguishing among linguistic operations that operate at distinct frequencies of neural oscillations (e.g., Bastiaansen and Hagoort, 2015). Low temporal resolution would also hinder the detection of differences in the timing at which different regions process the same incoming linguistic stimulus (Dehaene-Lambertz et al., 2006; Stephens et al., 2013; Uddén et al., 2019). In particular, given that regions across the core language network are anatomically connected and strongly synchronized in their activity patterns (e.g., Saur et al., 2008; Blank et al., 2014), information transfer across these regions is likely. Hence, by the time the fMRI BOLD signal is detected, an initially focal neural response might already appear ubiquitous throughout the network. We note, however, that whereas some studies using temporally sensitive methods report multiple, temporally and spatially separable profiles of linguistic integration (Zhang and Ding, 2017), others find that core language regions all exhibit highly similar responses (Fedorenko et al., 2016).

Beyond the methodological limitations of fMRI, studying the division of linguistic labor across the core language network also faces theoretical challenges. As discussed above, traditional distinctions between linguistic constructs (e.g., lexical semantics *vs.* combinatorial syntax) are no longer advocated by contemporary psycholinguistic theories—yet they still seem to guide a large portion of neuroimaging studies and neurobiological frameworks (e.g., Ullman, 2004; Friederici, 2012; Bornkessel-Schlesewsky and Schlesewsky, 2013; Friederici et al., 2017; for discussion, see Fedorenko et al., 2018). Moreover, which functional distinctions should be tested in lieu of the traditionally posited ones remains unclear; in the neuropsychological literature, despite strikingly different behavioral symptoms across some patients, the precise nature of these deficits in cognitive terms is still under debate (Caplan et al., 1996, 2007, 2013; Dronkers, 2000; Caramazza et al., 2001; Wilson and Saygin, 2004; Hillis, 2007; Grodzinsky and Santi, 2008). Perhaps, then, cognitive neuroscientific investigations of the division of linguistic labor across distinct mechanisms could better constrain cognitive models when their theoretical motivation is more grounded in current cognitive models and behavioral data.

### 4.3. A key methodology for neuroimaging studies of language processing

The current study tested whether a previously reported cortical hierarchy of integration timescales (Lerner et al., 2011) functionally corresponded to the core language network, and concluded that language regions all occupy a shared functional stage along this hierarchy. The apparent overlap between the spatial distribution of this hierarchy and the gross topography of the language network is therefore illusory. The key methodological innovation allowing us to demonstrate this point was augmenting the naturalistic paradigm for characterizing temporal receptive windows with a localizer task that identified core language regions in each individual brain. Such participant-specific functional localization established correspondence across brains based on response profiles rather than stereotaxic coordinates, thereby accounting for the substantial inter-individual variability in the precise mapping of function onto macro-anatomy (Amunts et al., 1999; Tomaiuolo et al., 1999; Juch et al., 2005; Caspers et al., 2006; Caspers et al., 2008; Scheperjans et al., 2008; Duffau, 2017). Without confounds in the data due to such variability, putative functional dissociations across temporal and inferior frontal regions dissolved, and a common integration timescale was established as a functional signature shared throughout the network.

When inter-subject correlations are instead computed using the common, anatomy-based approach (i.e., on a voxel-by-voxel basis), the resulting functional profiles at the group-level would often not be representative of any individual brain (compare Figures 3B and 2A). Interpretable ISCs would be obtained only in those stereotaxic coordinates that happened to consistently house the same functional unit across a sufficient number of participants in the sample. In contrast, in functionally heterogeneous cortical areas (Jones and Powell, 1970; Gloor, 1997; Wise et al., 2001; Chein et al., 2002; Fedorenko et al., 2012a; Deen et al., 2015), a single coordinate could belong to one functional network in some participants but to a second, distinct network in others (Poldrack, 2006; Fischl et al., 2008; Frost and Goebel, 2012; Tahmasebi et al., 2012), rendering signal “reliability” (and ISCs as a proxy thereof) an ill-defined concept. For instance, such distinct networks may reliably track respectively independent aspects of a given stimulus, resulting in low ISCs that do not adequately characterize either network. This issue pertains not only to studies of linguistic processing (e.g., Lerner et al., 2014), but to any study based on voxel-wise ISCs including, e.g., the many studies characterizing the episodic network (Honey et al., 2012b; Regev et al., 2013; Silbert et al., 2014; Simony et al., 2016; Baldassano et al., 2017; Lahnakoski et al., 2017; Yeshurun et al., 2017a; Yeshurun et al., 2017b; Regev et al., 2018), whose location is highly variable across individuals (Braga and Buckner, 2017).

Therefore, we urge researchers relying on ISC measures to confer functional interpretability to their findings by augmenting their approach with a methodology for establishing functional (rather than anatomical) correspondence across brains. For those who take issue with functional localizer tasks, alternative methodologies serving the same purpose are available (e.g., Haxby et al., 2011; Guntupalli et al., 2016). More generally, as naturalistic stimuli become a central tool in cognitive neuroscience due to their many advantages (Maguire, 2012; Ben-Yakov et al., 2012; Sonkusare et al., 2019; Wang et al., 2017; Hasson et al., 2010)—a trend that is celebrated in current volume—we should not do away with other, well tried approaches. Rather, a more promising way forward would be to synergistically harness the complementary strengths of multiple paradigms.

### 4.4. Conclusion

As linguistic inputs unfold over time, we integrate them into structured representations that mediate language comprehension. Whereas such integration might proceed hierarchically across the cortex, high-level language regions in the temporal and inferior frontal cortex all occupy a shared functional stage within this hierarchy. We find no evidence for a functional dissociation of integration timescales across these different regions of the core language network. Rather, they all exhibit sensitivity to information that extends from the morpheme- or-word level to the phrase- or sentence-level. This finding indicates that the division of linguistic labor across the core language network is not topographically organized according to the grain size of linguistic representation or by distinctions between more local *vs.* more global operations. Our results are instead more consistent with the notion of a spatially distributed set of tightly linked cognitive processes that integrate linguistic input into a continuum of increasingly larger constructions.

## 5. Acknowledgements

We acknowledge the Athinoula A. Martinos Imaging Center at the McGovern Institute for Brain Research, MIT. We thank Uri Hasson (Psychology Department and the Princeton Neuroscience Institute, Princeton University) for providing the stimuli as well as comments from himself and his colleagues on this work. E.F. was supported by a K99/R00 award HD 057522 from NICHD.

